# DNA methylation modulates transcription factor occupancy chiefly at sites of high intrinsic cell-type variability

**DOI:** 10.1101/003061

**Authors:** Matthew Maurano T, Hao Wang, Sam John, Anthony Shafer, Theresa Canfield, Kristen Lee, John A Stamatoyannopoulos

**Author notes:** These authors contributed equally to this work (Matthew Maurano T, Hao Wang).

## Abstract

The nuclear genome of every cell harbors millions of unoccupied transcription factor (TF) recognition sequences that harbor methylated cytosines. Although DNA methylation is commonly invoked as a repressive mechanism, the extent to which it actively silences specific TF occupancy sites is unknown. To define the role of DNA methylation in modulating TF binding, we quantified the effect of DNA methyltransferase abrogation on the occupancy patterns of a ubiquitous TF capable of autonomous binding to its target sites in chromatin (CTCF). Here we show that the vast majority of unoccupied, methylated CTCF recognition sequences remain unbound upon depletion of DNA methylation. Rather, methylation-regulated binding is restricted to a small fraction of elements that exhibit high intrinsic variability in CTCF occupancy across cell types. Our results suggest that DNA methylation is not a major groundskeeper of genomic transcription factor occupancy landscapes, but rather a specialized mechanism for stabilizing epigenetically labile sites.

## INTRODUCTION

DNA methylation is required for development and plays a central role in imprinting (Li et al. 1992; Jones 2012). Cytosine methylation in the context of CpG dinucleotides is also widely invoked as a causal mechanism for transcriptional repression at promoter regions (Jones and Taylor 1980). The correlation between DNA methylation and gene expression has long been recognized. Homozygous disruption of the DNA methyltransferases DNMT1 and DNMT3B in colorectal carcinoma cells (HCT116) stably reduces global DNA methylation by 83-95% (Rhee et al. 2002; Akalin et al. 2012), resulting in genome-wide changes in gene expression and chromatin structure (Pandiyan et al. 2013; Reddington et al. 2013). Chemical inhibition of DNA methyltransferases by 5-aza-2’-deoxycytidine (5-aza-CdR) transiently reduces global methylation levels and has been associated with reactivation of transcription factor binding or increased gene expression (Komashko and Farnham 2010; Qiu et al. 2010; Hagemann et al. 2011; Rodriguez et al. 2010; Witcher and Emerson 2009). However, to what extent such observed alterations in chromatin structure and gene expression are directly caused by demethylation of specific sites versus secondary effects is undetermined (Komashko and Farnham 2010; Bernstein et al. 2010; Stresemann et al. 2008; Yan et al. 2012; Tsai et al. 2012).

Recent findings indicate that dynamic demethylation during development is largely restricted to distal regulatory elements marked by DNase I hypersensitive sites (Hon et al. 2013). However, the mechanism(s) by which DNA methylation perturbs chromatin and regulatory mechanisms in a site-specific fashion remain obscure (Deaton and Bird 2011). A variety of possibilities for such effects have been suggested including potentiation of repressive chromatin features (Ramirez-Carrozzi et al. 2009; Collings et al. 2013), recruitment of methylcytosine-specific factors (Tate and Bird 1993; Lewis et al. 1992; Baubec et al. 2013), and direct interference with transcription factor (TF) occupancy (Tate and Bird 1993).

DNA methylation is specifically depleted at occupied transcription factor binding sites *in vivo* (Groudine and Conkin 1985; Lister et al. 2009; Neph et al. 2012b; Thurman et al. 2012). For example, the polyfunctional genome regulator CTCF exhibits a tight anticorrelation with DNA methylation at its binding sites *in vivo* (Phillips and Corces 2009; Wang et al. 2012), and its binding is abrogated by methylation *in vitro* (Bell and Felsenfeld 2000; Hark et al. 2000; Filippova et al. 2001; Renda et al. 2007). Although DNA methylation is widely assumed to inhibit TF occupancy, mechanistic studies suggest that methylation of TF recognition sites may follow TF evacuation (Brandeis et al. 1994; Lienert et al. 2011; Lin et al. 2000; Macleod et al. 1994; Matsuo et al. 1998; Stadler et al. 2011; Thurman et al. 2012; Feldmann et al. 2013). It nonetheless remains unclear whether altered methylation patterns themselves invoke transcriptional repression or are instead downstream consequences of other regulatory factors.

Here we investigate the causal relationship between genome-wide DNA methylation and TF occupancy using a model transcription factor (CTCF), whose binding is surmised to be regulated by DNA methylation (Phillips and Corces 2009). Using a combination of genetic and chemical depletion experiments coupled to genome-wide occupancy analysis, we find that the CTCF binding landscape remains largely unchanged in response to removal of DNA methylation. However, we observe a small minority of sites that reproducibly exhibit methylation-dependent occupancy. These sites are distinguished by high CpG content; the presence of specific CTCF recognition sequences that incorporate CpGs at critical positions in the binding interface; and an absence of preexisting regulatory activity within the assayed cell type. Despite the potential for CTCF occupancy at tens of thousands of potentially competent recognition elements genome-wide, reactivation is restricted specifically to elements that display highly variable CTCF occupancy when assayed across 40 cell types. Our results suggest that DNA methylation does not play a major role in repressing TF occupancy.

## RESULTS

### Limited reactivation of methylation-dependent sites

**Fig. 1.**
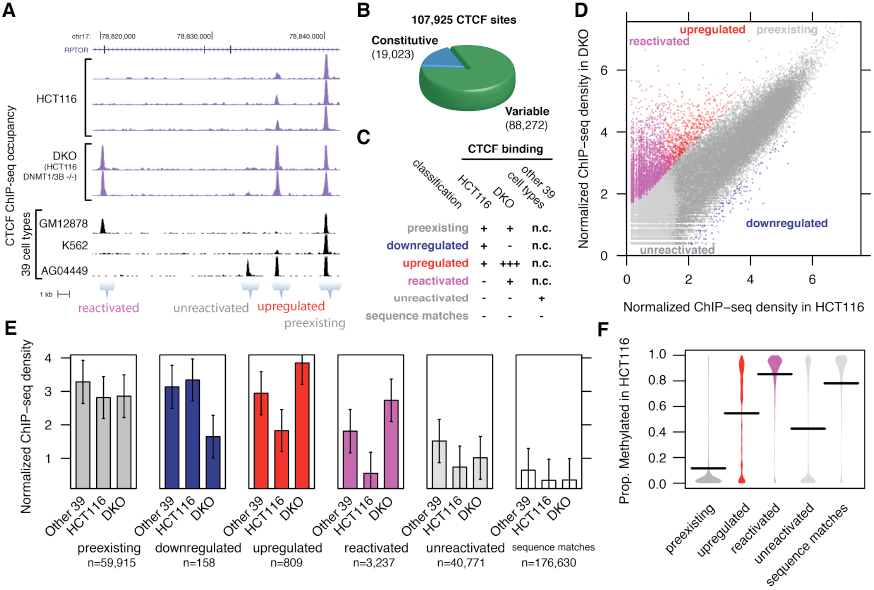
Profiling of methylation-silenced transcription factor binding sites in stably demethylated cells. (A) ChIP-seq profiling of CTCF occupancy in colorectal cancer cells depleted for methylation (HCT116, wild type; DKO, double knockout of DNMT1 and DNMT3B) shows reactivated sites at RPTOR locus present in DKO but not HCT116. (B) Profiling across 40 cell types comprehensively maps the variable CTCF *in vivo* binding landscape. (C) Classification scheme for CTCF sites based on ChIP-seq peak presence/absence (+/-) in HCT116; significant difference in occupancy upon demethylation in DKO (+ or +++, increased occupancy at 1% FDR); and peak presence/absence (n.c., not considered) in other 39 cell types. (D) A minority of sites exhibited increased occupancy in DKO (above diagonal). (E) Quantitative comparison of mean occupancy at each class of sites in HCT116, DKO or other cell types (measured as 90th percentile occupancy across 39 other cell types per site). Error bars represent standard deviation. (F) Average methylation across 9208 sites using RRBS profiling in HCT116. High methylation characterizes reactivated sites; unreactivated sites exist in both unmethylated and unmethylated populations. Downregulated sites (n=5 with methylation data) not shown. Horizontal bars represent mean.

We investigated CTCF occupancy in the context of reduced methylation by performing genome-wide profiling with chromatin immunoprecipitation (ChIP-seq) in HCT116 cells and DNMT1 and DNMT3B double knockout (DKO) HCT116 cells (Fig. 1A) including multiple biological replicates (3 for HCT116 and 2 for DKO). To measure the full range of cell type specificity at each CTCF site, we surveyed CTCF occupancy in 39 varied cell types, including new data for 20 cell types, each with at least two biological replicates (Table S1, Table S2). We identified a total of 107295 CTCF sites in these 40 cell types, DKO, and 5-aza-2’-deoxycytidine-treated K562 erythroleukemia cells (5-aza-CdR; described later). These data provide a comprehensive map of the binding patterns of the major genome regulator CTCF, representing a 38% increase in the number of known CTCF sites (Table S8). These data confirm that the CTCF binding landscape comprises a core minority of ubiquitous sites, but that the majority of its binding is cell-type specific (Fig. S1, Fig. 1B).

To precisely gauge the alteration to CTCF binding in the context of reduced genomic methylation, we categorized sites based on whether occupancy differed significantly (FDR 1%) between HCT116 and DKO and whether or not they were already occupied in HCT116 and (Fig. 1C and Methods). The majority of sites were ‘preexisting’ sites already bound in HCT116 with unchanged CTCF occupancy in DKO (Fig. 1D), suggesting a minor effect on a global scale. To confirm a lack of reactivation, we surveyed sequence matches genome-wide for the CTCF binding motif not exhibiting CTCF occupancy *in vivo* in any cell type (Fig. 1E, Fig. S2 and Methods). These sequences demonstrated a complete lack of reactivation, and exhibited a lack of occupancy similar to a set of random sequences without the CTCF recognition sequence. Thus the CTCF binding landscape remains largely similar despite a drastic reduction in genomic methylation.

However, we did identify a small compartment of 4204 sites that differed significantly in occupancy (FDR 1%), comprising 158 ‘downregulated’ and 809 ‘upregulated’ sites (bound in HCT116, but decreased or increased significantly); and 3237 ‘reactivated’ sites (DKO but not HCT116). Reactivated sites were strongly occupied, showing similar occupancy in DKO to preexisting sites in either cell type (Fig. 1E). To exclude the possibility that binding differences might be due to indirect effects from the targeting of the DNA methyltransferases rather than their subsequent effect on DNA methylation, we also profiled CTCF occupancy in single knockouts of both DNMT1 and DNMT3B with ChIP-seq. Each single knockout is known to exhibit little difference in methylation (Rhee et al. 2002), and correspondingly, we found that each deletion resulted in effectively no significant changes in occupancy (Fig. S3).

To verify whether reactivation was linked specifically to the relief of methylation-dependent repression, we examined a subset of CTCF sites for which quantitative profiling of methylation in HCT116 was available using reduced representation bisulfite sequencing (RRBS) (Varley et al. 2013). Confirming prior results (Lister et al. 2009; Wang et al. 2012; Thurman et al. 2012), methylation was low at preexisting sites and high at a majority of unbound sites (Fig. 1F and Table S3). Reactivation occurs almost exclusively (96%) at methylated sites. Yet only 50% of methylated unbound sites were reactivated, suggesting that alternate regulatory mechanisms prevent binding. Similarly, 90% of CTCF motifs without ChIP-seq occupancy were methylated but exhibited virtually no reactivation. Given the low global level of remaining methylation in DKO (Rhee et al. 2002; Akalin et al. 2012), it is unlikely the lack of reactivation represents localized persistence of methylation at unreactivated sites. These results suggest that while loss of methylation is necessary for CTCF binding at a subset of sites, methylation is not the sole barrier to *in vivo* occupancy at the majority of sites genome-wide.

### Reactivation of predetermined cell type-specific CTCF sites

**Fig. 2.**
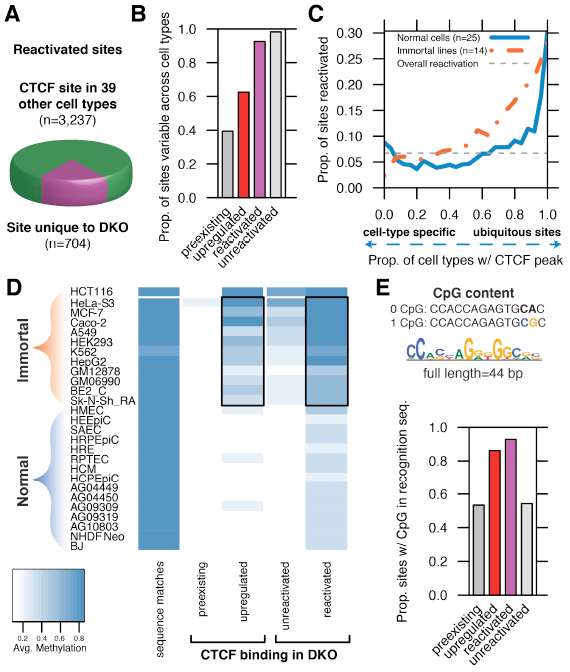
Reactivation of predetermined lineage-specific sites. (A) Most altered CTCF binding in DKO is at sites that vary in other cell types rather than novel binding to sequence matches. (B) Methylation-sensitive sites demonstrate high epigenetic instability across cell types (y axis; see Methods). (C) Higher reactivation frequency (y axis, reactivated sites as a proportion of both reactivated and reactivated sites) at sites found in all normal cell types but silenced in HCT116 (solid line, right), as well as immortal-only sites not found in normal lines (solid line, left). (D) Reactivated and upregulated sites are distinguished by increased methylation in immortalized lines (boxes). (E) The vast majority of reactivated and upregulated sites overlap a CpG in their recognition sequence.

We asked whether reactivated sites demonstrate CTCF binding in other cell types or if demethylation instead activates novel binding sites. Strikingly, we found that fully 91% of sites found in DKO but not HCT116 were found in at least 1 of 39 other cell types, suggesting reactivation occurs at a methylation-sensitive subset of sites with an inherent capacity for CTCF binding (Fig. 2A). However, there remain 44008 potentially reactivated sites found in other cell types but not HCT116, suggesting that methylation-sensitive sites are distinguished by specific characteristics. We use these potentially reactivated sites to compute the frequency of reactivation at various classes of CTCF sites.

By investigating CTCF binding across 39 other cell types, we found that upregulated and reactivated sites were far more likely than preexisting sites to vary epigenetically across cell types (Fig. 2B). Methylation sensitivity was highest at sites found in almost all cell types (excluding HCT116) and at sites found exclusively in malignancy-derived or immortalized lines (Fig. 2C). Methylation-dependent CTCF sites have been shown to regulate several tumor suppressors and oncogenes (Butcher et al. 2004; Witcher and Emerson 2009; Soto-Reyes and Recillas-Targa 2010; Dávalos-Salas et al. 2011), and a subset of methylation-associated cell-type selective CTCF sites exhibits characteristic hypermethylation in immortal but not normal lines (Wang et al. 2012), especially at CpG islands (Varley et al. 2013). We find that reactivated sites demonstrate the same pattern (Fig. 2D). The vast majority (93%) of reactivated CTCF sites had a CpG within the 44-bp region of protein-DNA interaction, compared to only 54% of unreactivated sites, and reactivation frequency increased with the quantity of CpGs at the binding sites (Fig. 2E). While 29% of the reactivated sites were in CpG islands, only 10% of unreactivated sites were. Thus, malignancy is associated with the disruption of a specific set of labile, methylation-sensitive CTCF sites.

### Chromatin dynamics at reactivated CTCF sites

**Fig. 3.**
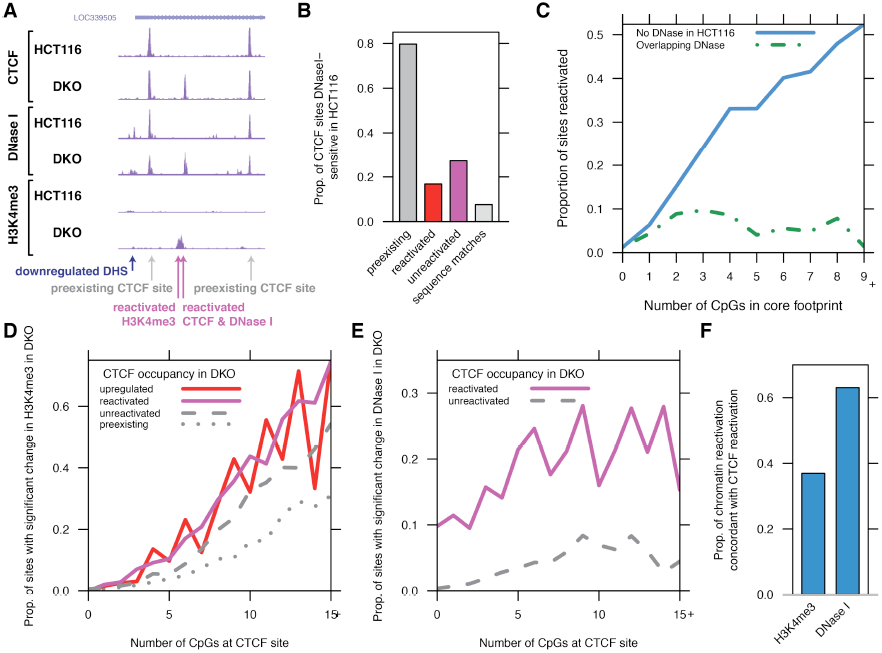
Chromatin dynamics at CTCF sites. (A) Profiling of CTCF, DNase I, and H3K4me3 reactivation in HCT116 and DKO. (B) Proportion of CTCF sites overlapping DNase I hypersensitive sites (DHSs). Although preexisting CTCF sites are stereotypically hypersensitive to DNase I cleavage, reactivation upon demethylation occurs at sites inaccessible in HCT116. (C) Reactivation frequency parallels local CpG content (solid line); while the presence of a preexisting DHS is associated with a near-total lack of reactivation regardless of CpG content (dashed line). (D-E) Increased CpG content (x axis) is strongly associated with reactivation of H3K4me3 (D) but not DNase I (E). Shown are sites without H3K4me3 peak (D) or DHS and CTCF peak (E) in HCT116. Note strong reactivation of H3K4me3 at unreactivated CTCF sites, in contrast to lack of increase in DNase I at these sites. (F) Reactivation of DNase I at CTCF sites coincides frequently with reactivation of CTCF occupancy, while H3K4me3 is more likely to increase without concordant CTCF reactivation.

The lack of reactivation outside of experimentally determined binding sites (particularly at methylated matches to the CTCF recognition sequence) implies that unoccupied CTCF sites are distinguished *in vivo* from a separate compartment of inert sequence matches. To examine whether chromatin accessibility prior to demethylation distinguishes these predetermined sites (John et al. 2011), we profiled HCT116 and DKO using DNaseI-seq and ChIP-seq for trimethylation of histone 3 lysine 4 (H3K4me3) and acetylation of histone 3 lysine 27 (H3K27ac), covalent modifications of histones associated with transcriptional activity and chromatin accessibility (Fig. 3A and Table S6). At the vast majority of CTCF sites, its occupancy is tightly associated with the presence of a DNase I hypersensitive site (DHS) marking increased chromatin accessibility (Fig. 3B). But, only 17% of reactivated sites overlapped DHSs in HCT116, suggesting that the novel recruitment of CTCF is not directed to chromatin accessible prior to demethylation (Fig. 3B).

In contrast to a lack of accessibility in HCT116 at reactivated CTCF sites, we observed that unreactivated CTCF sites were more frequently DHSs in HCT116 (Fig. 3B, Fig. S4A). Further confirming the lack of reactivation at sites of co-binding, the association of increased reactivation with increased number of CpGs was specific to sites without preexisting accessible chromatin (Fig. 3C). The presence of trimethylation of histone 3 lysine 4 (H3K4me3) was similarly antagonistic to reactivation (Fig. S4B), and the presence of both DNase I hypersensitivity and H3K4me3 was associated with a lack of methylation specifically at unreactivated sites despite a lack of CTCF occupancy (Fig. 1F, Fig. S4C). We found that unreactivated sites were also enriched for the occupancy of 17 transcription factors (Fig. S4D) studied by the ENCODE project (ENCODE Project Consortium 2012). Thus, sensitivity to methylation is reduced by the co-binding of additional regulatory factors permitting a relaxed chromatin state nevertheless inhospitable to CTCF binding.

Given that the removal of DNA methylation might indirectly enable CTCF occupancy subsequent to a broader relaxation of chromatin state (Ooi et al. 2007; Thomson et al. 2010), we then asked whether alterations in chromatin structure required the reactivation of CTCF binding. We found that DNase I accessibility increased in tandem with CTCF occupancy at reactivated, upregulated, and downregulated sites, but that only a subset of reactivated CTCF sites were accompanied by reactivation of H3K4me3 (Fig. S5). In fact, H3K4me3 reactivation occurred regardless of whether CTCF was recruited to the site and depended strongly upon CpG content (Fig. 3D), consistent with the observation that Cfp1 (part of the Set1 methyltransferase complex) recognizes unmethylated CpG islands (Thomson et al. 2010). In contrast, the correlation between DNase I reactivation and CpG content was almost entirely conditional on CTCF reactivation (Fig. 3E). While only a minority of H3K4me3 reactivation was associated with CTCF reactivation, most increased DNase I accessibility was at sites with CTCF reactivation (Fig. 3F). Thus, we conclude that while active histone marks such as H3K4me3 are deposited in regions of high CpG content regardless of CTCF occupancy, DNase I accessibility specifically marks regulatory factor occupancy.

The extent to which other mammalian TFs interact with the global methylation landscape remains obscure (Lienert et al. 2011). Thus to contextualize the contribution of CTCF in connecting DNA methylation to regulatory sites genome-wide, we examined the full accessible chromatin landscapes of HCT116 and DKO cells using DNaseI-seq. DKO cells showed a generally expanded accessible chromatin landscape, comprising a 61% increase in the number of DHSs (Fig. S6), but this represents a surprisingly small increase given the even greater reduction of DNA methylation. We found that the vast majority of H3K4me3 (84%) and H3K27ac (88%) peaks in DKO overlapped with DHSs, confirming the global sensitivity of DNase I profiling for regulatory activity. Using a stringent threshold, we found 12279 DHSs with statistically significant differential accessibility (1% FDR), 96% of which were reactivated DHSs not present in HCT116. We found that 70% of reactivated DHSs were present in a survey of 124 cell types (Maurano et al. 2012a), confirming preferential reactivation of silence sites from other lineages. We found that between 48% (1% FDR) and 77% (5% FDR) of reactivated CTCF sites without a DHS in HCT116 were recognizable from chromatin accessibility alone, suggesting a high sensitivity to reactivation of individual factors. Although CTCF has a large number of binding sites and plays a pivotal role at imprinted loci, 90% of reactivated DHSs were not CTCF sites, suggesting a broad potential pool of methylation-responsive regulatory sites (Fig. S6D).

### CpGs at key positions in the protein-DNA binding interface drive reactivation

Although increased CpG content is strongly associated with CTCF reactivation (Fig. 3C), it is unclear whether this represents a nonspecific relaxation of high-CpG regions upon removal of DNA methylation or a specific disruption of the CTCF protein-DNA interface. Consistent with a direct effect on CTCF binding, we observed a strongly increased reactivation (up to 27%) at the best matches to the CTCF consensus sequence (Fig. S7A). We classified CTCF binding sites by sequence similarity into three known binding modes (Filippova et al. 1996; Ohlsson et al. 2001; Bowers et al. 2009; Rhee and Pugh 2011; Nakahashi et al. 2013), including engagement of zinc fingers 3-7 (core sites) and additional specific interactions by zinc fingers 8-10 (upstream and extended spacing sites). We found that core sites were reactivated 42% more frequently than upstream sites (Fig. S7B), supporting speculation that different modes of engagement of its 11 zinc fingers may confer susceptibility to different modes of regulation or functional selectivity (Ohlsson et al. 2010). The markedly increased reactivation frequency for core sites may represent increased sensitivity to CpG methylation due to the lower binding energy from the reduced protein-DNA binding interface. Despite the correlation of sequence features with reactivation, it is important to reiterate that local sequence content alone is insufficient to distinguish *in vivo* occupied sites from other recognition sequences in the genome (Fig. S2C).

**Fig. 4.**
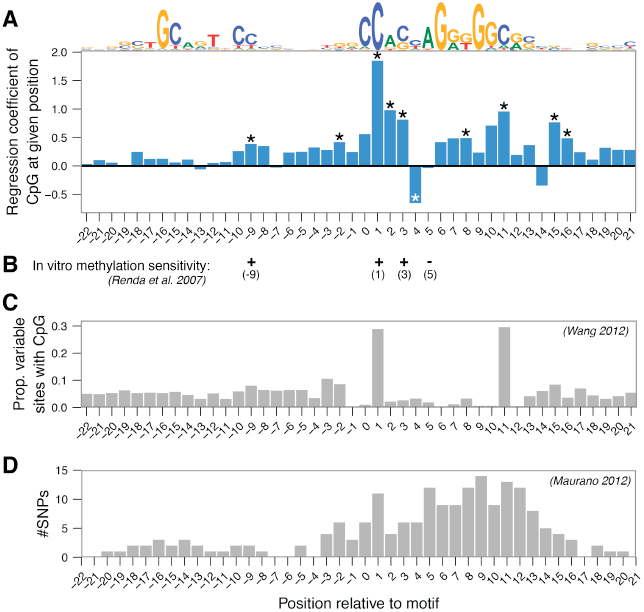
CpGs at key positions in the protein-DNA binding interface drive reactivation. (A) Linear regression estimates (y axis) of the contribution of the presence of a CpG dinucleotide at each position (x axis) in CTCF recognition sequence (consensus binding motif shown at top) to reactivation upon demethylation. Window defined as the 44 bp extent of protein-DNA contact demarcated by DNase I footprint. *, Position is significant in regression (P < 0.05, Bonferroni-corrected). (B) Methylation-sensitive positions identified *in vitro* through electrophoretic mobility shift assay (EMSA) (Renda et al. 2007). (C) Association of CpG presence with methylation-associated cell-type variability across 19 cell types (Wang et al. 2012) (Fig. 4). (D) Single nucleotide polymorphisms (SNPs) associated with significant alteration in occupancy across individuals (Maurano et al. 2012b) (Fig. 3B).

To examine whether the association of CpG content with reactivation localized to specific positions in the protein-DNA interface (Renda et al. 2007; Wiench et al. 2011; Wang et al. 2012), we used a linear regression model to quantify the contributions of CpG dinucleotides in the recognition sequence. We found that critical CpGs were concentrated in the core binding region, consistent with *in vitro* sensitivity to methylation (Renda et al. 2007), the positions of CpGs at sites of methylation-associated cell-type selective binding (Wang et al. 2012), and sensitivity to single nucleotide variants (Maurano et al. 2012b) (Fig. 4). Although it is possible that different CTCF binding modes exhibit differential sensitivity to methylation at specific locations within the recognition sequence, we did not find a significant effect of binding mode on susceptibility at any given position.

### Cell type-specific reactivation highlights methylation-independent repression

**Fig. 5.**
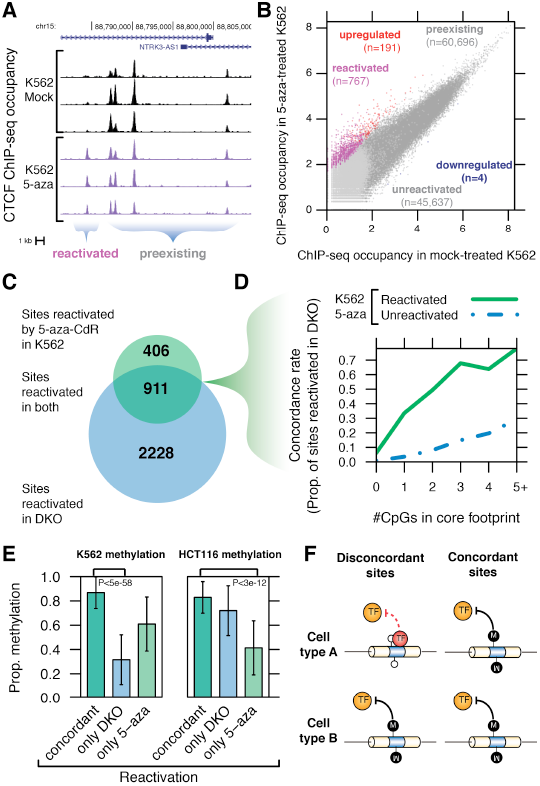
Cell type-specific reactivation highlights methylation-independent repression. (A) Reactivation at NTRK3 locus showing reactivated and preexisting sites in a mock-and 5-aza-CdR-treated K562 cells. (B) Limited but reproducible reactivation (FDR 1%). Sites were classified (labels) based on presence of peak in mock-treated K562 and significant differential occupancy in treated cells. (C) Majority of 5-aza-CdR-reactivated sites are concordantly reactivated in DKO (of a potential 46,726 sites not occupied in both K562 and HCT116, FDR 5%). (D) Sites reactivated in K562 (solid line) are more likely to be reactivated in DKO (y axis) than unreactivated sites (dashed line) regardless of CpG content. (E) Reduced preexisting methylation (y axis) in both K562 (left) and HCT116 (right) distinguishes sites reactivated specifically by genetic or chemical means, suggesting that non-concordant sites are attributable to methylation-independent silencing. P-values by Wilcoxon signed-rank test. (F) Concordant reactivation implies methylation-dependent silencing across cell types, while discordant reactivation implies methylation-independent silencing in the unreactivated cell type.

To test whether the compartment of methylation-sensitive CTCF sites was consistently reactivated in a system using transient depletion of DNA methylation, we performed ChIP-seq on mock-treated and 5-aza-CdR-treated K562 cells. We observed weaker reactivation with 5-aza-CdR than in DKO cells, including 767 reactivated, 191 upregulated, and 4 downregulated sites, DKO (Fig. 5A-B). We assessed the extent of demethylation with high resolution using a capture sodium bisulfite design (Methods, Table S7). Consistent with previous reports (Rhee et al. 2002; Hagemann et al. 2011; Pandiyan et al. 2013), these results confirm a more limited and uneven reduction in methylation than the almost completely demethylated DKO cells (Fig. S8). CTCF reactivation was largely specific to sites of preexisting methylation: 79% of reactivated sites were methylated (>25%) in K562 cells. However, only 30% of reactivated sites showed altered methylation (>5% difference) and 38% of unreactivated sites showed no methylation change, suggesting widespread secondary effects and a similarly muted response of CTCF binding to demethylation. Conversely, only 12% of the CTCF sites methylated and unoccupied in K562 exhibited reduced methylation (>5%) upon treatment (Fig. S8D). This complex relationship is consistent with previous observations that 20% of variable CTCF sites are unmethylated in all cell types (Wang et al. 2012), and that most 5-aza-CdR-induced alterations in gene expression or chromatin structure occur at previously unmethylated sites (Komashko and Farnham 2010).

Despite the lesser extent of reactivation with 5-aza-CdR, we found substantial overlap of reactivation: 69% of sites reactivated with 5-aza-CdR overlapped sites also reactivated in DKO (Fig. 5C). Sites reactivated concordantly in both cell types demonstrated CpG-dependent reactivation (Fig. 5D), and exhibited high preexisting methylation in both K562 and HCT116 cells, further supporting the existence of a predetermined class of methylation-dependent sites. We sought to specifically identify sites reactivated in only one cell type upon depletion of methylation, as a given unoccupied CTCF site might be silenced in a different fashion between K562 and HCT116. Indeed, sites discordantly reactivated in only one of the two cell types demonstrated significantly less methylation (Fig. 5E), suggesting the presence of a methylation-independent silencing mechanism in the unreactivated cell type (Fig. 5F).

### Recognition of labile methylation-sensitive sites

**Fig. 6.**
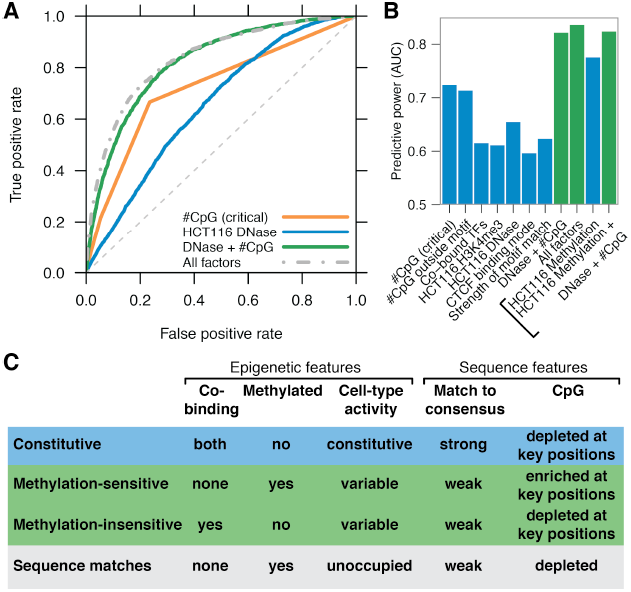
Recognition of labile methylation-sensitive sites. (A) Predictive power of features to distinguish reactivated from unreactivated sites, illustrated by receiver-operator curves (ROC). Area under the curve (AUC) summarizes overall predictive power. Dashed gray line indicates a random predictor and has an AUC of 0.5. A perfect predictor would be plotted as a right angle. (B) Detailed predictive power (y axis) of selected genomic features. Models including methylation (bracketed) measured only at sites with RRBS data. Green bars indicate linear models combining multiple factors. Other TF occupancy refers to the sum of HCT116 ENCODE TF ChIP-seq track densities. (C) CTCF sites exhibit distinct classes of sensitivity to methylation.

Given the importance of methylation at regulatory sites in imprinting and malignancy, we examined the predictive power of genomic characteristics to recognize the minority of sites that were reactivated in comparison to the unaffected CTCF binding landscape (Fig. 6). The number of CpGs at the critical positions in the recognition sequence (Fig. 4A) was the single most predictive factor (Fig. 6A). The number of CpGs in the flanks was almost as predictive, probably due to the generally high CpG density of CTCF sites (Fig. 6B). The second most predictive factor was lack of chromatin accessibility in HCT116 (Fig. 6A). As many of the features surveyed are not independent, we used a logistic regression model to assess the combined predictive power of the characteristics we identified (Methods). A model considering just chromatin accessibility and number of CpGs was more predictive (AUC of 0.82) than either factor alone, but almost as predictive as the full model (AUC 0.84), indicating that the remaining factors provide only minor additional discriminative power (Fig. 6B). Although we observed increased reactivation at exons, the increased G+C content of protein-coding sequence is presumably responsible for a markedly increased reactivation frequency relative to other genomic sequence (Fig. S9A), which suggests an altered regime of methylation-regulatory factor interaction at these sites (Stergachis et al. 2013). At the subset of sites for which methylation data were available, the degree of methylation alone had strong predictive power (Fig. 6B). But this predictive power was largely redundant when DNase I accessibility and number of CpGs was additionally considered, suggesting that the latter features primarily provide indication of methylation status (Fig. S4C). Thus, reactivated sites can be reliably distinguished based on *in vivo* profiling and a simple model considering CpG content and chromatin accessibility (Fig. 6C).

## DISCUSSION

We describe a surprisingly muted response of the regulatory landscape to large-scale depletion of methylation, using both transient and stable inhibition of DNA methyltransferases in two different cell types. Although some sites showed exquisite sensitivity to methylation, demonstrating increased occupancy despite a small changes in methylation, the binding landscape as a whole remains mostly unchanged. This is consistent with the moderate effect on binding of single nucleotide changes in the CTCF recognition sequence (Maurano et al. 2012b), and suggests that its rich binding interface is buffered at most sites *in vivo* to the effect of methylation. Given the global lack of correlation between demethylation and altered binding, these results place a limit on the extent of CTCF-mediated coupling between DNA methylation and genome organization. Thus although DNA methylation globally shows a tight relationship with reduced CTCF binding, this relationship is only causal at a small subset of sites, cautioning against a facile interpretation of alterations of DNA methylation in oncogenesis.

That reactivation is strictly limited to *in vivo* CTCF sites from other cell types suggests that either these silent sites are globally marked in HCT116 or that other sequence matches to the CTCF recognition sequence are silenced independent of methylation. Approximately half of both reactivated and unreactivated sites are accessible to DNase I in human embryonic stem cells, compared to only 11% of sequence matches, suggesting that the marking of bona fide CTCF sites occurs early in development. But it remains unclear how unreactivated sites are distinguished from mere sequence matches. These silent sites are not marked by DNase I accessibility, and in fact reactivation appears to be less frequent in the presence of cofactors. Thus, although CTCF preferentially recognizes a DNA sequence with higher information content than most other human transcription factors, it appears likely that cooperative binding with as-yet undetermined factors plays an even larger role in the determination of its *in vivo* binding sites.

Despite the secondary role of methylation at most sites, a set of reproducibly reactivated sites is distinguished by solitary CTCF binding and clear sequence characteristics including the presence of CpGs at key positions in the binding interface. The strongest factor distinguishing reactivated sites is the presence of CpG dinucleotides at key positions of the protein-DNA recognition interface. This localization is consistent with our previous findings showing that the depletion of methylation is specifically increased at sites of transcription factor occupancy marked by DNase I footprints (Neph et al. 2012b). The conferral of methylation-sensitive binding by the presence of CpG dinucleotides at key positions is reflected in epigenetically lability across cell types and suggests that TF binding provides a key link between sequence features, DNA methylation, and epigenetic state (Neph et al. 2012b; Gaidatzis et al. 2014).

## MATERIALS AND METHODS

### Cell culture and ChIP-seq/DNaseI-seq

HCT116 and DKO cells have been previously described (Rhee et al. 2002). Cells were cultured in appropriate medium and maintained in a humidified incubator at 37°C in the presence of 5% CO2. (Table S1). K562 cells were treated with 5-aza-2’-deoxycytidine (decitabine; Sigma, A3656) dissolved in DMSO to 10mM. Drug was administered at 1uM daily for 3 days. Control K562 cells were mock-treated with DMSO. Conditions for other cell types as described (Table S1).

Chromatin immunoprecipitations were performed as described for CTCF (Wang et al. 2012) and H3K4me3 and H3K27ac (Thurman et al. 2012). Briefly, cells were crosslinked for 10 min in 1% formaldehyde and quenched in 125 mM glycine. Chromatin was sheared by Bioruptor (Diagenode) and incubated with antibody conjugated to Dynabeads (M-280, Invitrogen). After reversing crosslinks, immunoprecipitated DNA was purified by phenol-chloroform-isoamyl alcohol extraction and ethanol precipitation. DNase was performed as described (John et al. 2011; Thurman et al. 2012), whereby small (300-1000 bp) fragments are isolated from lysed nuclei following DNase treatment. Libraries generated from immunoprecipitated or DNase-treated DNA were sequenced on an Illumina Genome Analyzer IIx or HiSeq 2000 by the High-Throughput Genomics Center (University of Washington) according to a standard protocol. Most experiments were performed on two independent biological replicates (Table S2, Table S4, Table S5).

### Data processing

ChIP-seq and DNase data were mapped to the human genome (GRCh37/ hg19) using bowtie (Langmead et al. 2009) with the options ‘‘bowtie -n 2 -m 1 -e 70 --best’’ allowing up to three mismatches. Reads mapping to multiple locations were then excluded, and reads with identical 5’ ends and strand were presumed to be PCR duplicates and were excluded using Picard MarkDuplicates. Smoothed density tracks were generated using bedmap (http://code.google.com/p/bedops/) to count the number of tags overlapping a sliding 150-bp window, with a resolution of 20 bp (Neph et al. 2012a). Density tracks for display were normalized for sequencing depth by a global linear scaling to 10 million tags.

CTCF data from 39 cell types was processed as previously (Wang et al. 2012). Briefly, a master peak list was established from ENCODE project 2% IDR-processed SPP peak calls (Kharchenko et al. 2008; Li et al. 2011) (ftp://ftp.ebi.ac.uk/pub/databases/ensembl/encode/supplementary/integration_data_jan2011/byDataType/peaks/june2012/spp/optimal/), with locations adjusted to center on matches to the nearest CTCF motif (P < 10-4, fimo) if the motif was within 50 bp. Peak presence in a given cell type was established by the presence of a 1% FDR hotspot (John et al. 2011) (also http://www.uwencode.org/proj/hotspot/) in any replicate. Peaks only found in DKO and/or 5-aza-CdR-treated K562 cells were not included in total peak counts to maximize their generality. We identified possible further CTCF binding sequences using the program fimo (P<10^−5^), as well as random sequences without a CTCF binding sequence (P>10^−4^). Both sets were required to be further than 500 bp from any known *in vivo* CTCF site, to consist of base pairs 90% of which are uniquely mappable by 36 bp reads, and not to be on the ENCODE blacklist (ENCODE Project Consortium 2012). We identified DNase I hypersensitive sites by the presence of a hotspot (FDR 1%) in any HCT116 replicate or a hotspot (FDR 0.5%) in DKO; novel sites in DKO were included in the analysis if they overlapped no unthresholded hotspot in any HCT116 replicate. H3K4me3 and H3K27ac regions were called using hotspot.

At both CTCF sites and DHSs separately, we measured CTCF occupancy or DNase I accessibility by the number of tags overlapping the 150-bp region, and H3K4me3 and H3K27ac density by tags overlapping the 2150-bp region. We then used the package DESeq (Anders and Huber 2010) to identify significant differences. CTCF ChIP-seq peaks and DHSs discordant between the two previously published replicates and the new replicate for HCT116 (FDR 10%) were removed. Pairwise differences were called between samples for each ChIP-seq and DNase I data set using estimateDispersions(method="pooled", sharingMode="maximum", fitType="local") and nbinomTest. After observing a near-complete lack of reactivation at sequence matches and random sites (<10 sites), we subsequently excluded these sites and recomputed pairwise differences. We applied the Benjamini-Hochberg procedure to control for multiple testing. To obtain normalized data at all sites, we used estimateDispersions as before but with fitType=”parametric”, and then applied a variance stabilizing transformation and averaged the occupancy of all replicates. These values were scaled to [0, 10] by subtracting the global minimum and dividing by the global maximum * 10.

We obtained location of CTCF sites relative to genes using RefSeq and relative to CpG islands and repetitive regions in RepeatMasker from the UCSC Genome Browser.

### Classification of CTCF motif models

CTCF binds in a multivalent fashion, whereby three modes of binding are distinguished by the presence and position of an upstream motif (Nakahashi et al. 2013). At each site we chose the best-matching (by fimo P-value) of three motif models.

### Per-nt regression model

We used the lm() function in R to perform a binomial regression considering all preexisting, upregulated, unreactivated and reactivated CTCF sites with a recognizable motif:

reactivation ∼ c + x-22 + ... + x21 + upstream + (#CpGs in flanking region)

Upstream represents an indicator variable for the presence of either upstream motif. Multiple testing correction was performed using the Bonferroni method.

ROC and PR curves were computed using the R package ROCR. Binomial models for multiple factors were constructed using the lm() function in R.

### Monitoring methylation

Purified DNA from mock-and 5-aza-CdR-treated K562 cells was fragmented following the Agilent SureSelect Methyl-Seq protocol with slight modifications (Table S7). 4 ug from each of the samples was fragmented in a Covaris S2 (Covaris) under the following conditions: 10% duty cycle at intensity 5 for 60 seconds with 200 cycles per burst for 6 cycles with mode set to sweeping. The 250 bp fragmented DNA was then end repaired and adenylated followed by ligation to adapters synthesized with 5’ –methylcytosine instead of cytosine. The adaptor-ligated library was purified using AMPure XP beads (Beckman Coulter Genomics). 500 ng of each library was hybridized to Agilent SureSelect Methyl-Seq biotinylated RNA baits (84 Mbp) for 24 h at 65 °C. The biotinylated probe/target hybrids were captured on Dynal MyOne Streptavidin T1 (Invitrogen), washed, eluted, and desalted following purification on a MinElute PCR column (Qiagen) as described in the SureSelect protocol.

Bisulfite conversion of the purified captured library was performed using the EZ DNA Methylation Gold Kit (Zymo Research) as per manufacturer’s instructions. The bisulfite converted captured library was amplified by PCR with minimal amount of PCR cycles, then purified using AMPure XP beads and quantified by Qubit dsDNA assay (Invitrogen) following the SureSelect protocol.

Samples were diluted to a working concentration of 10 nm. Cluster generation was performed for each sample and loaded on to single lane of an Illumina HiSeq flowcell. Single end sequencing was performed for 36 cycles according to manufacturer’s instructions.

Capture bisulfite reads were mapped to the human genome (hg19) using Bismark (Krueger and Andrews 2011) and bowtie2 beta 6 (Langmead et al. 2009) with the options “-n 1” and excluding duplicate reads and combining reads from the top and bottom strands. Methylation was monitored at CpGs with ≥8 read coverage, and averaged for all CpGs within the 150 bp window at each CTCF site. We obtained ≥8x coverage in both samples for 658,010 CpGs.

RRBS data for 29 cell types is available from the UCSC Genome Browser (Varley et al. 2013), processed as described (Wang et al. 2012).

### Data access

Data have been submitted to the NCBI Gene Expression Omnibus (GEO) (http://www.ncbi.nlm.nih.gov/geo/) under accession nos. GSE30263 and GSE50610 (http://www.ncbi.nlm.nih.gov/geo/query/acc.cgi?token=hjypvqoommwekjo&acc=GSE50610).

Data sets are also available for viewing in the UCSC Genome Browser (http://genome.ucsc.edu/).

## ACKNOWLEDGEMENTS

We thank Bert Vogelstein for graciously sharing HCT116 DNMT1/3B DKO cells. We thank Bob Thurman for informatics support. This work was supported by NIH fellowship F31MH094073 to M.T.M. and grant U54HG004592 to J.A.S..

